# An end-to-end pipeline based on open source deep learning tools for reliable analysis of complex 3D images of Medaka ovaries

**DOI:** 10.1101/2022.08.03.502611

**Authors:** Manon Lesage, Jérôme Bugeon, Manon Thomas, Thierry Pécot, Violette Thermes

## Abstract

Computational analysis of bio-images by deep learning (DL) algorithms has made exceptional progress in recent years and has become much more accessible to non-specialists with the development of ready-to-use tools. The study of oogenesis mechanisms and female reproductive success in fish has also recently benefited from the development of efficient three-dimensional (3D) imaging protocols on entire ovaries. Such large datasets have a great potential for the generation of new quantitative data on oogenesis but are, however, complex to analyze due to imperfect fluorescent signals and the lack of efficient image analysis workflows. Here, we applied two open-source DL tools, Noise2Void and Cellpose, to analyze the oocyte content of medaka ovaries at larvae and adult stages. These tools were integrated into end-to-end analysis pipelines that include image pre-processing, cell segmentation, and image post-processing to filter and combine labels. Our pipelines thus provide effective solutions to accurately segment complex 3D images of entire ovaries with either irregular fluorescent staining or low autofluorescence signal. In the future, these pipelines will be applicable to extensive cellular phenotyping in fish for developmental or toxicology studies.

**Summary statement:** An accessible image analysis method for biologists, which includes easy-to-use deep learning algorithms, designed for accurate quantitative measurement of ovarian content from complex 3D fluorescent images.

## INTRODUCTION

As imaging methods for thick biological samples improve and become more widespread in various fields of life sciences, the volume of image data keeps growing and their analysis becomes even more complex. Biologists are therefore facing a rising need for computational tools to analyze large bio-image datasets and extract reproducible and meaningful biological information.

The fish ovary is a complex organ that shows important structural and functional changes during reproductive cycles. It contains different types of cells, including oocytes (*i*.*e*., female gametes) and numerous surrounding somatic supporting cells that form, together with each oocyte, the functional units known as ovarian follicles (Lubzens et al., 2010; Nakamura et al., 2009). During oogenesis, each follicle grows and differentiates until finally giving rise to eggs that are ultimately released during spawning. One of the greatest challenges facing research on the development of ovarian dynamics and functions is the lack of an effective method to accurately count growing oocytes regardless of their stage. Studies have indeed traditionally been limited to automatic or manual oocyte counting on two-dimensional (2D) ovarian sections and extrapolation of the data to the whole organ or to manual counting of dissociated follicles (Gay et al., 2018; Iwamatsu, Takashi, 1978; Iwamatsu, 2015). Some studies have also focused on the development of complex stereological approaches to limit the biases induced by 2D approaches (Charleston et al., 2007). Recently, the emergence of optical tissue clearing methods and powerful microscopes have opened new perspectives with the possibility of imaging whole ovaries in three dimensions (3D), notably for mice and fishes (Fiorentino et al., 2021; Lesage et al., 2020; Soygur and Laird, 2021). It is thus now possible to generate 3D image data, generally of very large size, that ideally allows direct and comprehensive access to all structures and to achieve a precise 3D image reconstruction of the whole ovary. However, tools for 3D image analyses are still too inaccurate and tedious, especially for image segmentation, partly because of an irregular contrast signal in depth and the presence of oocytes of heterogenous sizes, as reported previously for the adult Medaka ovary (Lesage et al., 2020). Ovarian 3D imaging therefore has a promising future, but its widespread use still relies on the availability of more efficient and easier-to-use computerized analytical tools.

In recent years, artificial intelligence (AI) has developed considerably and is proving to be highly effective for digital image analysis in biology, which has recently led to a deluge of publications in this field. Various algorithms based on deep learning (DL) have emerged and provide many applications in microscopy allowing to overcome classical limitations such as image segmentation. They allow to increase object recognition accuracy, segmentation reproducibility and enable to save a considerable amount of time for the analysis of large datasets by limiting manual interventions of users (Moen et al., 2019). Some specific methods have thus been proposed to automatically segment follicles in the mammalian ovary from histological 2D sections using a convolutional neural network (CNN) (İnik et al., 2019; Sonigo et al., 2018). Other more generalist tools have recently emerged to democratize the use of DL technology with few prerequisites in computed coding, by providing either DL trained models accessible from public databases (https://bioimage.io/#/), notebooks accessible from any computer (von Chamier et al., 2021), or other open-source plugins such as CSBDeep (Weigert et al., 2018) or DeepImageJ (Gómez-de-Mariscal et al., 2021). Among the available models for cell segmentation, Cellpose is a particularly versatile one, providing a generalist pre-trained model for segmentation that can perform on various cell types in a great variety of acquisition modalities (Stringer et al., 2021). Cellpose has recently proven to be very effective in segmenting muscle fibers from 2D images of histological sections (Waisman et al., 2021). Noise2Void (N2V) is another approach that stands out for its image denoising performance. N2V does not require noisy image pairs nor clean target images, therefore allowing training directly on the corpus of data to be denoised (Krull et al., 2019). In the era of deep learning, it thus appears that some of the routine limitations for bio-image analysis are now solved. All that remains for the biologist is the delicate task of integrating deep learning steps into the various analytical procedures for 2D and for 3D images in particular.

The aim of this study was to test the possibility of using a pre-trained open-source model to improve the critical step of segmentation of Medaka ovary 3D images without undergoing the fastidious and complex task of neural network training. We generated 3D fluorescent images of the adult ovarian follicle boundaries, by using the Methyl Green nuclear dye. We also generated 3D images of ovaries at the larvae stage, by using the autofluorescence signal in oocyte cytoplasm. For 3D segmentation of both types of images, we applied the generalist Cellpose model for oocyte 3D segmentation, which was even more efficient after image pre-processing steps and N2V denoising. A post-processing step after Cellpose was also set up to eliminate any remaining error and to combine labels when necessary. N2V and Cellpose have thus been integrated into a complete pipeline that allows an accurate estimation of the oocyte content from complex 3D images of the whole Medaka ovary.

## MATERIAL AND METHODS

### Ethical Statement

All fish were reared in the INRAE ISC-LPGP fish facility, which hold full approval for animal experimentation (C35-238-6). All fish were handled in strict accordance with French and European policies and guidelines of the INRAE LPGP Institutional Animal Care and Use Committee (no.M-2020-126-VT-ML, no.M-2019-48-VT-SG).

### Medaka breeding and sample collection

Medaka fish (*Oryzias latipes*) from the CAB strain were raised at 26°C under artificial photoperiod dedicated to growth phase (16 h light/ 8 h dark) or reproductive cycles (14 h light/ 10 h dark). Female fish were sampled either at larvae stage (20 days post-hatch, dph) or adult stage (5 months old). Fish were euthanized by immersion in a lethal dose of MS-222 at 300mg/L supplemented with NaHCo_3_ at 600mg/L and fixed overnight at 4°C in 4% paraformaldehyde (PFA) diluted in 0.01 M phosphate buffer saline (PBS) pH 7.4. Larvae were then dehydrated gradually in methanol and stored at -20°C. Adult ovaries were dissected after fixation and directly stored at 4°C in PBS + 0.5% (w/v) sodium azide (S2002, Sigma-Aldrich).

### Fluorescent staining and clearing

Larvae were progressively rehydrated in PBS and ovaries were dissected. Ovaries were then permeabilized and immunostained following the iDISCO protocol with some modifications (Renier et al., 2014). Samples were successively incubated in PBS/0.2% Triton X-100 (PBSTx) for 30 min twice, PBSTx/20% DMSO for 30 min at 37°C and in PBSTx/0.1% Tween-20/20% DMSO/0.1% deoxycholate/0.1% NP40 at 37 °C for 3 h. Ovaries were washed in PBSTx for 15 min twice, then blocked in PBS/0.1% Triton X-100/20% DMSO/6% Sheep Serum for 2H30-3H at 37°C. Samples were immunolabelled with anti-phospho-Histone H3 (Ser10) primary antibody (1:500, 06-570 Merck millipore), washed for 0.5 day in PBS/0.1% Tween-20/10μg/ml heparin (PBSTwH) under gentle agitation, and incubated with Alexa-Fluor 546 secondary antibody (1:500, A11035, ThermoFisher). Antibodies incubations were conducted for 2.5 days at 37°C in PBSTwH/5% DMSO/3% Sheep serum. Finally, stained larvae ovaries were embedded in low-melting agarose 1% before proceeding to clearing. Adult ovary samples were stained and cleared according to the C-ECi method with few modifications (Lesage et al., 2020). For staining, adult ovaries were incubated with the Methyl Green dye (MG) (40 μg/mL, 323829, Sigma-Aldrich) in PBS/0.1% Triton X-100 at 37°C for 2.5 days. After Staining, both adult ovaries and embedded larvae ovaries were dehydrated in serial methanol/H2O dilution series supplemented with Tween-20 (2% and 0.1%, respectively), then immersed in 100% ethyl-3-phenylprop-2-enoate (ethyl cinnamate [ECi]) (W243000, Sigma-Aldrich) and finally kept at room temperature until subsequent imaging step.

### Samples mounting and imaging

Image acquisitions were performed with a Leica TCS SP8 laser scanning confocal microscope equipped with a 16x/0.6 IMM CORR VISIR HC FLUOTAR objective (ref. 15506533, Leica, Wetzlar, Germany). For larvae ovaries, samples embedded in agarose blocks were glued on a coverslip and placed in a glass Petri dish filled with ECi. Adult ovaries were successively placed with ventral side up or down for complete imaging despite the objective working distance limitation, and mounted as described previously (Lesage et al., 2020). Mosaic z-stack tiles were stitched in Leica software using 11,72% overlap. Larvae ovaries were acquired in 1024×1024 pixels, 400Hz (unidirectional) with an optical zoom of 1.3 and a z-step of 1.63 μm (voxel size 0.52 × 0.52 × 1.6264 μm). PH3 fluorescent signal was acquired using 552 nm laser excitation slightly above optimal intensity (3-4%), and frame average was set to 2. Acquisitions took between 1.5 and 5.5 hrs according to ovary size and generated 1 to 2 GB of data. Adult ovaries were acquired in 512×512 pixels, 600Hz (bidirectional), optical zoom 0.75, z-steps 6 μm (voxel size 1.80 × 1.80 × 6.00 μm), line accumulation 2 and frame average 2. Ventral and dorsal z-stacks were acquired in about 10 hrs each and generated 8 to 10 GB of data. MG staining was detected with 638 nm laser and excitation gain compensation was used along Z axis (5 to 10% intensity).

### Image processing

A schematic overview of image treatment workflows is shown on Figure 1. All steps were conducted on the open-source FIJI software, unless otherwise specified.

**Figure 1:**
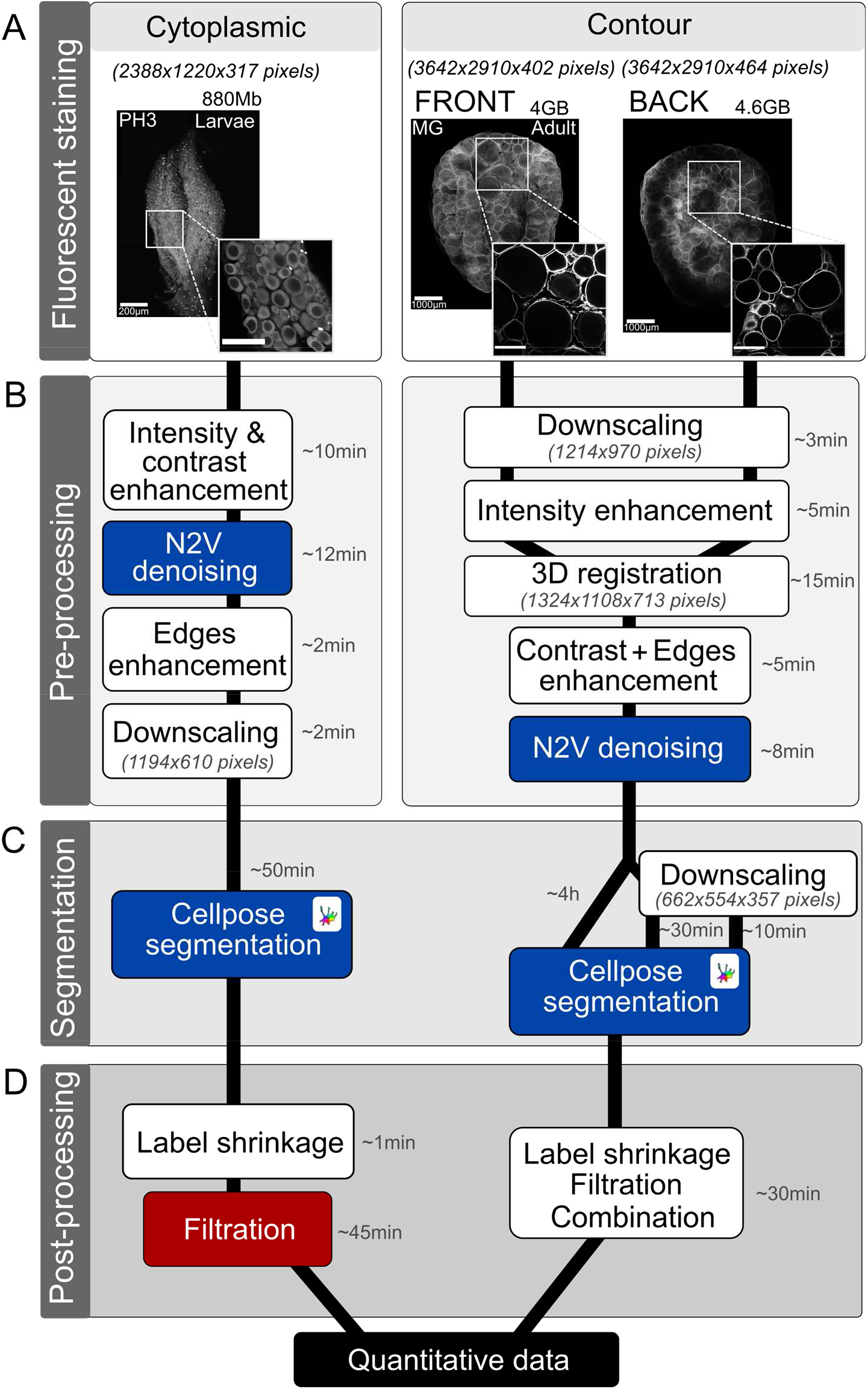
Pipeline overview for 3D image analysis of the whole ovary at larvae and adult stages. (A) Fluorescent staining strategies for whole ovary imaging. Cytoplasmic autofluorescence from Histone H3 phosphorylation immunofluorescence (PH3) is used for larvae (left panel). Methyl-green (MG) nuclear staining delineating follicles contour is used for adult stage (right panel). Z-projection of raw data are shown (standard deviation method. Raw stacks sizes are indicated. (B) Image pre-processing steps used for reconstruction and enhancement, listed from top to bottom. (C) 3D segmentation step is performed with Cellpose algorithm. Adult images are subjected to several segmentations runs before and after image downscaling. (D) Image post-processing is performed for segmentation correction, label filtering and final quantitative analysis. Opensource tools are indicated in white boxes, deep-learning opensource tools in blue, commercially available software in red (AMIRA). Relative computation time for one sample is indicated (B-D). Voxel size is indicated in brackets (A-C). Scale bars 200 μm (for larvae), 1000 μm (for adult), 100 μm (for larvae inset), 500 μm (for adult inset)

#### Image intensity and contrast enhancement

Before image enhancement, adult z-stacks were downscaled in order to reduce computation time. A resampling factor of 3 on X and Y axes was used, which resulted in images sizes of 1214 × 970 pixels. Progressive intensity and gamma correction plugin was applied along the Z axis to compensate fluorescence loss in depth (Fig. 1B). For larvae, exponential or linear interpolation method were used with default parameters and intensity enhancement was set between 150 and 400% depending on samples. For adults, linear interpolation method was used, intensity set between 200 and 800% and normalization was selected (modifying range of pixel intensity values by linear scaling method). A linear gamma correction was also performed (factor 1.5) to enhance mid tones pixels on adult images. Image contrast was then enhanced by applying Contrast Limited Adaptive Histogram Equalization (CLAHE) with following parameters: block size 128, bins 256, slope 3 and fast mode, for larva; block size 512, bins 256, slope 30 and fast mode, for adult. A Fiji macro was used to apply this function on Z-stacks by batches, which is available on our GitHub page: https://github.com/INRAE-LPGP/ImageAnalysis_CombineLabels.

#### 3D registration

Adult ventral and dorsal 3D stacks were registered, aligned and combined with the Fijiyama plugin using the “two images registration mode (training)” (Fig. 1B). A manual registration was first performed to roughly superimpose the two volumes. Automatic registration was then applied for linear image transformation with block-matching alignment method. Linear transformations included rigid transformations (translation and rotation) and, if necessary, similarities transformations (rigid and isotropic homothetic factor). The two registered stacks were fused with Image calculator (Max operator), resulting in image size of 1324 × 1108 × 713 pixels.

#### Signal-to-noise ratios enhancement

Three-dimensional images were denoised using Noise2Void (N2V) deep-learning based tool available on Fiji, using a model trained on a few selected 3D stacks snippets. A 2D model was trained on folder containing ∼15 Z-stacks snippets (512×512, from 50 to 115 z-steps) cropped from different larvae samples. Training patch shape was set at 96×96 pixels and N2V automatically used data augmentation (90, 180 and 270 rotations and flipping), thereby multiplying total patches amount by 8. The resulting pool of 2D patches were used for training (90%) and validation (10%). Training was performed with 250 epochs, 150 steps/epoch and batch size set to 128, resulting in ∼13 hrs of training with our computer specifications. Denoising prediction duration was estimated to ∼12 min for 1GB of data with our stated parameters (batch size 2). For adult, similar strategy was used for training, using 10 z-stack snippets (256×256, 100-200 Z-steps), patch shape 64×64 pixels. Training was performed with 300 epochs, 200 steps/epoch, batch size 128, for a total of ∼9 hrs of training. Denoising prediction duration was estimated to ∼8 min for 1GB of data with our hardware specifications and stated parameters (batch size 2). For image edges enhancement, stacks were subjected to a 3D median filter (x,y,z radius 1,1,1 for larvae and 2,2,2 for adult). Filtered image was then subjected to external morphological gradient computation (shape: ball; x,y,z radius 3,3,3 for larvae and 2,2,2 for adult) with Morphological filters (3D) function of MorpholibJ plugin. External gradient image was then subtracted from original pre-treated stack (without median filtering). An internal morphological gradient was also computed on larvae stacks (element shape: ball, x,y,z radius 4,4,4) and added to image data. For 3D visualization of data, volume reconstructions were performed on the Amira software using Volren rendering (Fig 2A, E) or Volume-rendering (Fig 3A, F and 4A, F).

**Figure 2:**
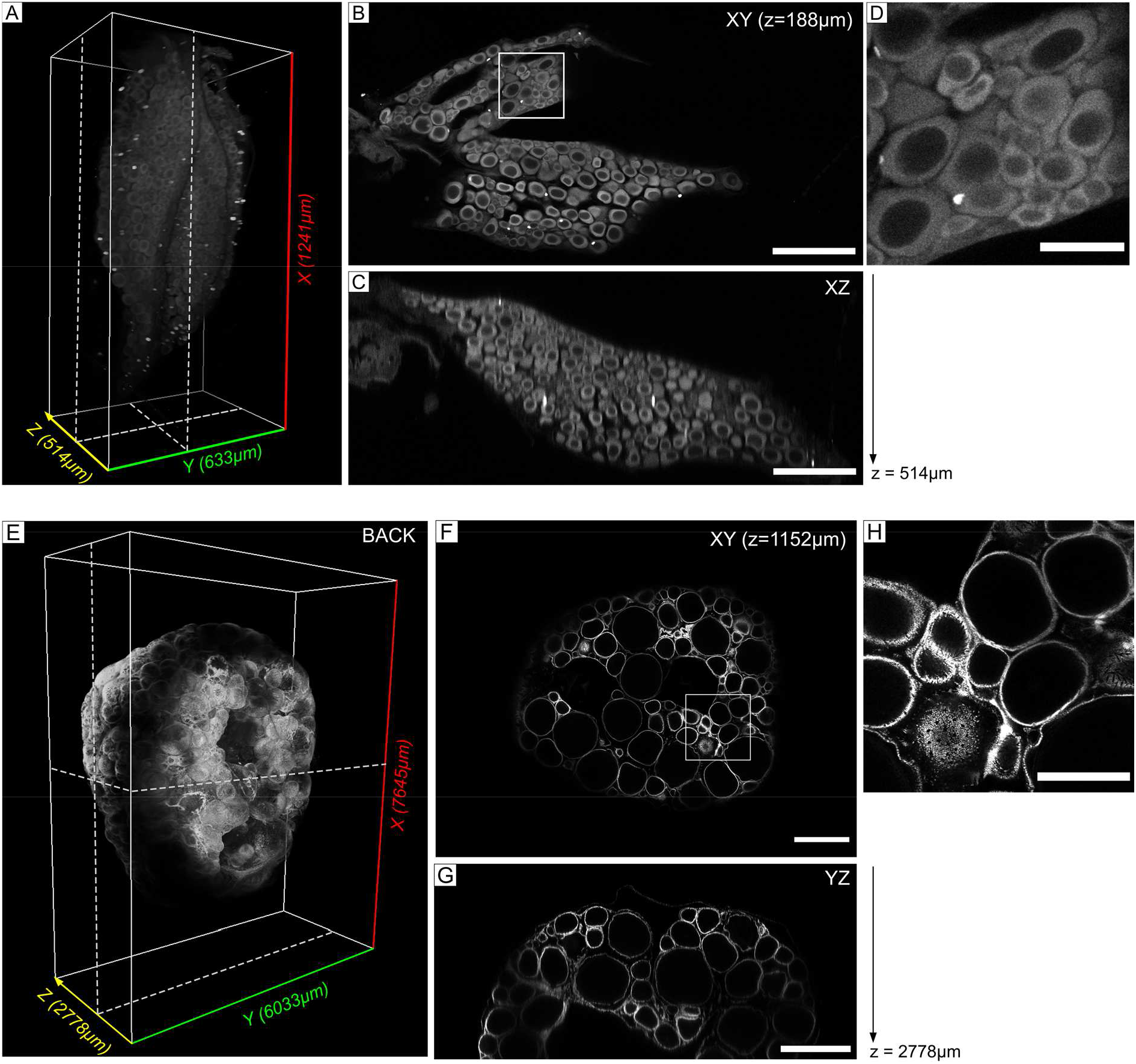
3D reconstruction of whole medaka ovaries. (A) Larvae ovary reconstruction with raw data. Total ovary size approach 1240 μm in length (x), 630 μm in width (y) and 515 μm in height (z). XY plane at 188μm and XZ plane are shown with dotted lines. (B) XY plane showing PH3 staining and cytoplasmic background in oocytes at 188μm in depth, magnified in inset (D). (C) XZ orthoslice of larvae PH3 staining. A decrease in fluorescence intensity is observable near 400 μm in depth. (E) Adult ovary reconstruction with raw data. Only back stack (dorsal face) is shown, with a size of 7645 μm in length (x), 6033 μm in width (y) and 2778 μm in height (z). XY and YZ virtual slices are shown with dotted lines. (F) XY plane at 1152μm depth shows MG staining resulting in delimitated follicular contours, magnified in inset (H). (G) YZ orthoslice of dorsal face of adult ovary. Heterogeneity of MG staining is observable through depth. Scale bars 200μm (B, C), 50μm (D), 1000μm (F, G), 40μm (H).

**Figure 3:**
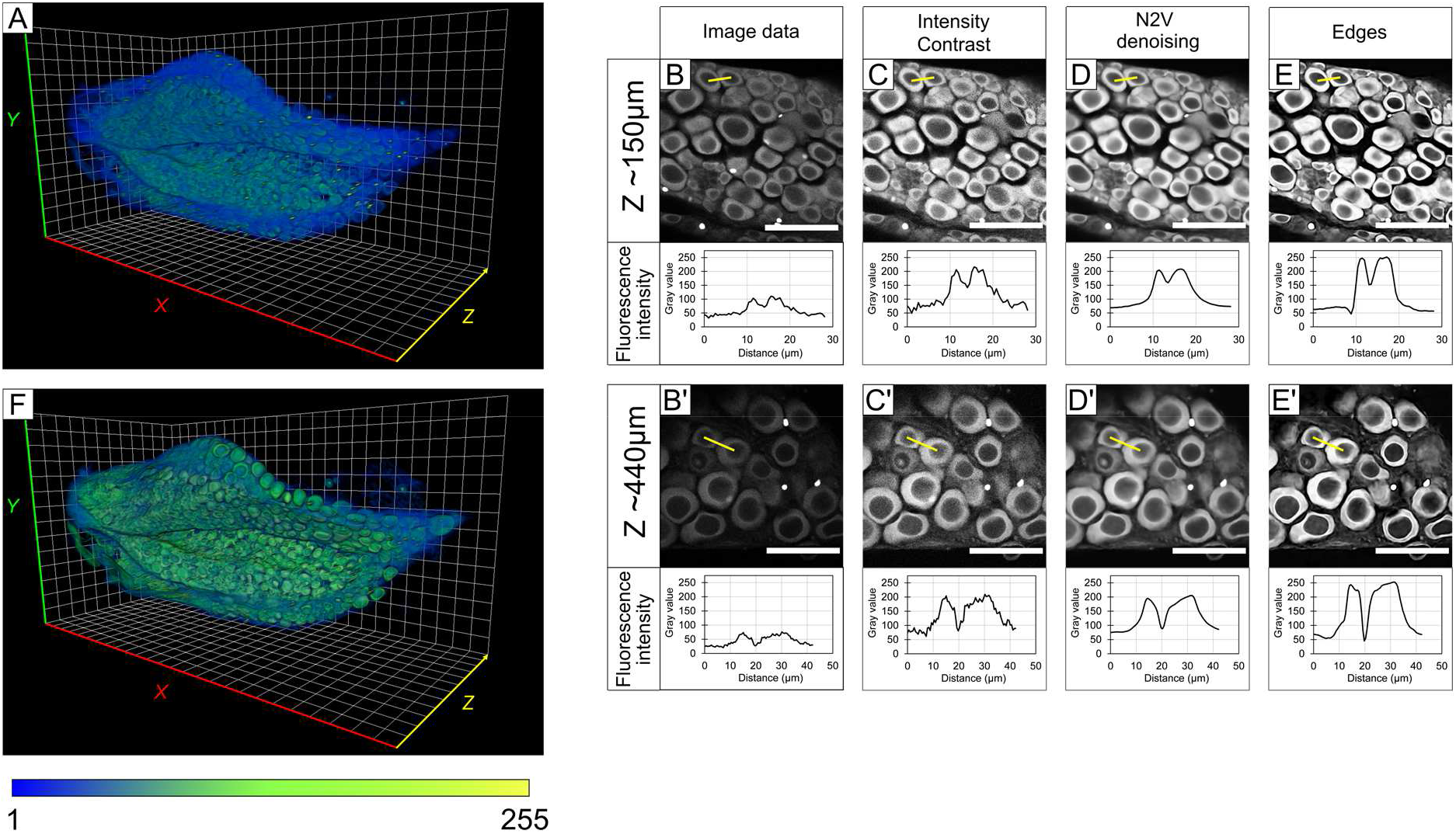
Image pre-processing for enhancement of features detection through larva ovary depth. (A) Representative 3D reconstruction of 20 dph (days post-hatching) larvae ovary before image processing. (B-E) Effect of successive image processing steps at 150 μm depth and (B’-E’) at 400 μm depth assessed on XY cropped planes. A profile line intensity is used to assess fluorescence intensities nearby relevant objects to be segmented. Fluorescence intensity and signal to noise ratio are progressively enhanced. (F) 3D reconstruction of 20 dph larvae after image pre-processing showing signal homogenization. Color gradient is representative of grey levels (1-255). Scale bar 100 μm, Grid square size 50 μm.

**Figure 4:**
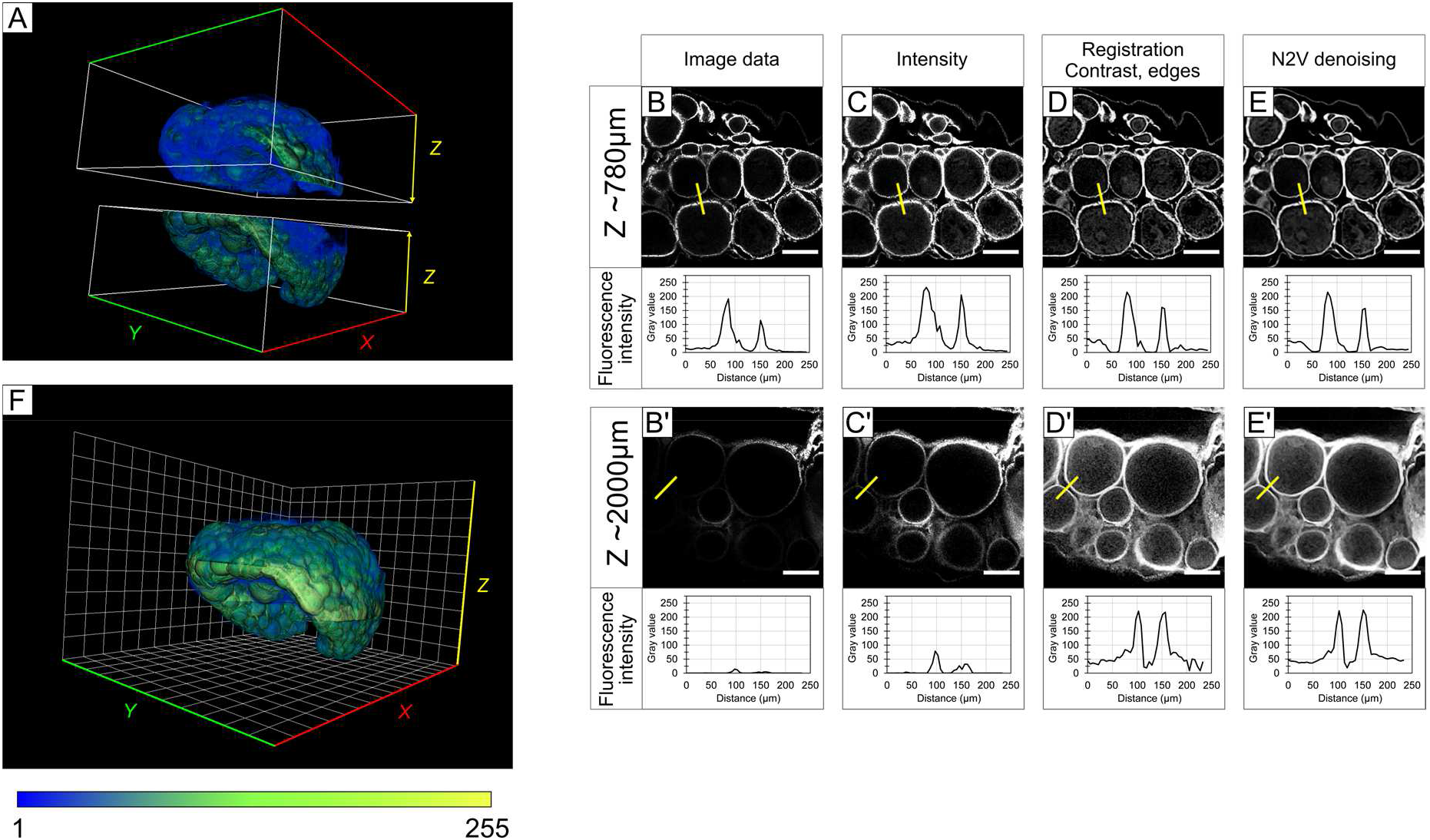
Image pre-processing for enhancement of features detection through adult ovary depth. (A) Volume reconstruction of front and back adult z-stacks before image processing. (B-E) Effect of successive image processing steps at 780 μm deep and (B’-E’) at 2000 μm deep assessed on XY cropped planes extracted on front stack. A profile line intensity is used to assess fluorescence intensities nearby relevant objects to be segmented. Fluorescence intensity loss in depth is greatly recovered and resolution of follicles contours is improved. (L) Final adult ovary reconstruction after 3D registration and image enhancements. Color gradient representative of grey levels. Scale bar 300 μm, Grid square size 500 μm.

#### Deep learning 3D segmentation

Follicle segmentation was performed using Cellpose algorithm with local environment installation, launched from Anaconda command prompt (Fig. 1C). For larvae, X and Y scale were first reduced by half so that mean follicle diameter approach ∼30 pixels, which is the optimum diameter for Cellpose cell segmentation (final image size 1194 × 610 pixels). Cellpose was then run in 3D with “cyto” pre-trained model, setting parameters as follows: diameter 30, cellprob threshold -2, anisotropy 1,6, min size 10. A batch size of 2 was used, depending on GPU memory allocation, resulting in ∼50 min for segmentation prediction of ∼250 Mb of data. Resulting masks were saved in TIFF format for subsequent data treatment. For adults, the same process was used except anisotropy was set to 1.7. 3D segmentation took ∼4 hrs for ∼1 Gb of data. To segment out-of-range follicles, adult stacks were downscaled once more by applying a resampling factor of 2 in X, Y and Z (no interpolation, final images size of 662 × 554 × 357 pixels). Downscaled stacks were subjected to Cellpose segmentation with diameter size set to 30 and 60 pixels. 3D segmentation took ∼35 min and ∼11 min for ∼125 Mb of downsized data, for 30 and 60 pixel diameter respectively.

#### Post-processing and data extraction

For post-processing of segmented follicles, data were first slightly narrowed. For that operation, label boundaries were computed with MorpholibJ plugin and subtracted from the original Cellpose results. For larvae ovary images, labels were then post-processed on AMIRA software. Labels were subjected to an opening morphological operator (3 pixels, precise) and then filtered based on their size (Equivalent Diameter >= 1.5e-5m) and shape (ShapeVAa3d <=3.5). Few remaining errors were manually corrected. For adult ovary images, label shrinkage, filtration and combination were performed automatically or semi-automatically using a Fiji macro. Labels were filtered based on volume and sphericity parameters. When necessary, segmentation images were rescaled to match 3D registered image size (1324 × 1108 × 713px). The combination strategy consisted in adding largest segmented labels from downscaled images (using Cellpose diameter 30) on original scale label segmentation (Cellpose diameter 30) where largest follicles were over-segmented (Supplemental Fig. 1). Briefly, labels >650μm volume-equivalent diameter (EqDiameter) were filtered from the downscaled image and added to the original scale label image after selection and deletion (using morphological reconstruction operation) of wrong labels resulting from over-segmentation. For combination of missing largest labels, a similar strategy was used with downscaled label image obtained with Cellpose diameter 60, but with semi-automatic method. Missing labels were manually selected with multi-point tool and then processed as presented before. Macro “CombineLabels” was developed in IJ1 Macro language and can be downloaded from the Github page: https://github.com/INRAE-LPGP/ImageAnalysis_CombineLabels. The volumes of all segmented follicles were exported and equivalent diameters were calculated. For adults, EqDiameter were subjected to a correction factor of 1.12 to compensate the volume shrinkage due to sample clearing, as described in Lesage *et al*. (Lesage et al., 2020). Data analysis was performed on labels above 25 and 50 μm in diameter, for larvae and adult samples respectively. Label 3D reconstructions were generated on Amira using volume-rendering object.

### Hardware and software

Data were analyzed on a 64-bit Windows 10 Pro computer equipped with a 2x Intel Xeon Silver 4110 (8 Cores, 3.0GHz) processor, a Nvidia Geforce GTX 1080 graphic card, and 384 Go of RAM. We used the Amira 2020.2 software with the XLVolume extension (Thermo Fisher Scientific, Waltham, Massachusetts, United States), Anaconda3-2021.11 python distribution, Python 3.7.9, CUDA toolkit 10.0, PyTorch 1.6.0 and Cellpose v0.6.1 (Stringer et al., 2021). We also used FIJI (Schindelin et al., 2012) and the following plugins: CLAHE (Pizer et al., 1987; Zuiderveld, 1994), Progressive intensity and gamma correction (Murtin, 2016), Fijiyama (fijiyama-4.0.0) (Fernandez and Moisy, 2021), CLIJ2 (clij2-2.5.3.0, Haase et al., 2020), MorpholibJ (morpholibJ-1.4.3, Legland et al., 2016), Noise2Void (n2v-0.8.6)(Krull et al., 2019) and CSBDeep (csbdeep-0.6.0)(Weigert et al., 2018).

## RESULTS

### 3D imaging of the ovaries

To detect oocytes within the ovary at both adult and larvae stages, sample were fluorescently stained and optically cleared to allow full imaging by confocal fluorescence microscopy (Fig. 1A). For adult ovaries, nuclei of supporting cells surrounding the oocytes were stained with the fluorescent nuclear dye Methyl-Green (MG) identified as a convenient marker for delineating follicle boundaries (Lesage et al., 2020). For larvae ovaries (20 dph), which are composed of small early developing oocytes flanked by only a few supporting somatic cells, we took advantage of the cytoplasmic autofluorescence generated by immunostaining (here anti-phospho-histone H3 antibody, PH3). Resulting images displayed a very low signal-to-noise ratio (SNR) and a rapid loss of signal recovery in depth for larvae ovaries (Fig. 2A-D). Signal intensity was twice as low at 440 μm in depth compared to the top (150 μm depth, Fig. 3B, B’). In addition, it is noteworthy that smaller oocytes were less distinguishable than larger ones having thicker cytoplasm, especially in very compact regions (Fig. 2D and 3B). Images stacks of adult ovaries displayed a higher fluorescence signal with a high SNR that was recovered up to 1152 μm in depth, although some heterogeneity in fluorescence intensity was observable (Fig. 2F, G). At a greater depth (2 000 μm), images display a substantial loss of signal intensity (Fig. 4B, B’).

### Image enhancement and 3D visualization

Given the uneven signal intensity of the images, and especially the very low SNR observed with the non-specific staining of larval ovaries, we applied successive processing steps to enhance the fluorescent signal throughout Z-stacks prior segmentation. For larvae ovary, fluorescence intensity of image stacks was progressively enhanced along Z-axis to increase the signal in depth, and mean grey values were increased and homogenized to enhance contrasts (Fig. 3B, B’, C, C’). To minimize the noise potentially introduced by intensity and contrasts adjustment and to avoid potential aberrant enhancement of noisy structures, image stacks were denoised using the self-supervised N2V deep-learning-based algorithm (Krull et al., 2019) (Fig. 3D, D’). Finally, edges were refined using morphological gradients (Fig. 3E, E’). XY views from Z-stacks and fluorescent intensity profiles through adjacent oocytes show the progressive signal recovery over the different steps at both 150 and 440 μm in depth. It is noticeable that while normalizing grey values distribution, N2V denoising preserves oocytes edges with limited blurring effect, thus minimizing any feature loss (Fig. 3D, D’). In addition, it is noteworthy that overexposure was created in some cases as a side effect of edge refinement. The challenge here was therefore to find a compromise between the loss of detection of underexposed oocytes and the overexposure generated in order to achieve the greatest difference between light and dark levels. Image pre-processing steps thus enabled to increase the overall fluorescent signal intensity, to better define edges of the oocytes and to homogenize the fluorescence intensity across the Z-stack, thereby allowing a better 3D reconstruction of the larval ovary (Fig. 3A, F).

For 3D images of adult ovaries, a similar strategy was applied except an extra step of automatic 3D registration that was performed for the reconstruction of the whole ovary (Fig. 4B-E and 4B’-E’). As a result of the combination of images in the overlap region, 3D registration led to a slight increase in fluorescence intensity in this region in the final stack (Fig. 4A, F). XY views of Z-stacks and fluorescent intensity profiles through adjacent oocytes showed a significant increase of the SNR, especially at 2 000 μm in depth. Similar to the larvae ovary, the pre-processing allowed to improve the fluorescence signal, and especially to homogenize the fluorescence intensity through the Z-stack for a better 3D reconstruction of the adult ovary (Fig. 4A, F).

### Cellpose efficiently identifies oocytes and follicles on 3D images

For 3D oocyte and follicle segmentation on larvae and adult images, we selected the open-source Cellpose deep-learning algorithm because of its generalist nature for cell segmentation (Stringer et al., 2021). We compared the efficiency of Cellpose for 3D segmentation before and after image pre-treatment. In both cases, Cellpose could detect either internal fluorescent staining (oocyte cytoplasm) or external fluorescent staining (somatic follicular cells), on larvae and adult ovary images respectively (Fig. 5A-D and 5E-H). Notably, Cellpose was much more efficient on pre-treated images than raw images. Although XZ views of larvae stacks revealed accurate segmentation along the Z axis, several undetected oocytes and some Z-label fusions were detectable in the absence of preprocessing (Fig. 5B and 5D, insets). For adult ovaries, segmentation of raw images leads to many cases of over-segmentation in conjunctive tissues or in large follicles, as well as fewer detected follicles, compared to segmentation of pre-processed images (Fig. 5F and 5H, insets).

**Figure 5:**
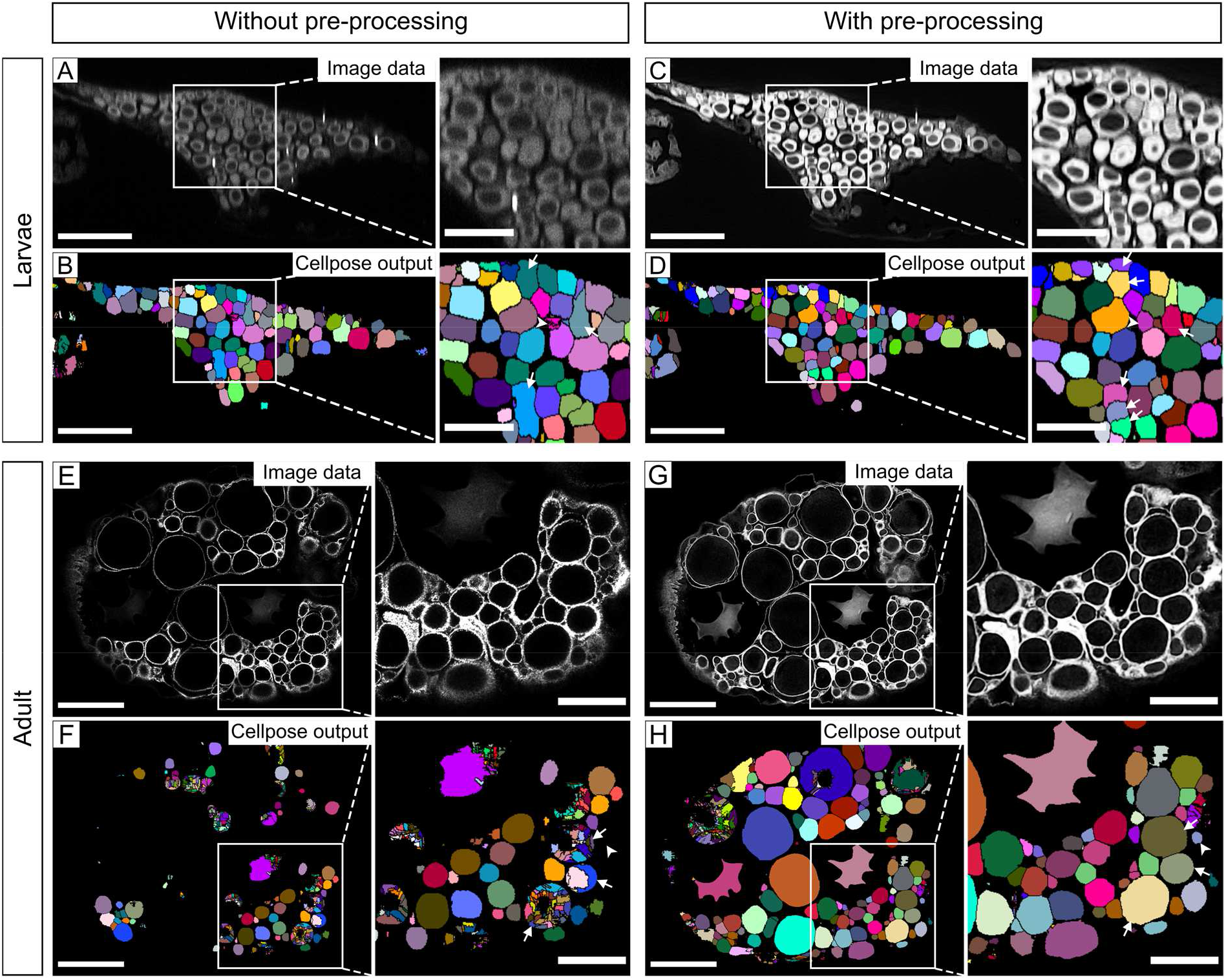
Effect of image pre-processing on Cellpose 3D segmentation efficiency. (A) XZ orthoslice of larvae ovary showing raw data and (C) image data after pre-processing, magnified on insets. (B) Cellpose segmentation output using raw data as input or (D) pre-processed image data. Results are shown after label erosion to correctly visualize labels individualization. Insets show more segmentation errors without image pre-processing, including unsegregated labels (arrows) or over-segmentation (arrowhead). (E) XY plane of adult ovary on raw data and (G) after image processing, magnified on insets. (F) Cellpose segmentation output for 30 pixel diameter using either raw data input or (H) pre-processed image. Segmentation results show high error number without image pre-processing, including many missing labels (arrowhead) especially in locations with heterogenous staining and high over-segmentation in medium and large size follicles. Scale bars: 200 μm (A-D), 100 μm (insets in A-D), 1000 μm (E-H), 500 μm (insets in E-H).

### Post-processing of label images after Cellpose 3D segmentation

Cellpose output images were post-processed to adjust the label sizes to that of the oocytes (label shrinkage) and to remove outliers (label filtration) (Fig. 6). Label shrinkage was performed by automatically subtracting the label boundaries to the original Cellpose labels. For adult ovaries that have the unique feature of containing heterogeneous follicle sizes (ranging from about 20 to 1 000 μm in diameter), different Cellpose label images were generated by modulating image resolution of the input image (Fig.6B, C). If necessary, a 60 pixel diameter was used for Cellpose segmentation to detect largest follicles. The different resulting label images were combined in an additional post-processing step by using a Fiji Macro named “CombineLabels” (Fig. 6D). Images of larvae ovary labels show that, after post-processing, the majority of labels perfectly fit to the shape and size of the oocytes and that aberrant labels with elongated shapes or very small sizes were removed. In few cases, some inaccuracies still persisted, mainly under-segmentation of small oocyte clusters (Fig. 6A, arrows) or non-segmented oocytes (Fig. 6A, arrowhead). Similar to larvae ovary images, results of segmentation and post-processing of adult ovary images were highly accurate, both in terms of follicle detection, label shape and size fitting (Fig. 6B-D). After post-processing, remaining segmentation errors were limited to a few outlier labels located outside the relevant structures.

**Figure 6:**
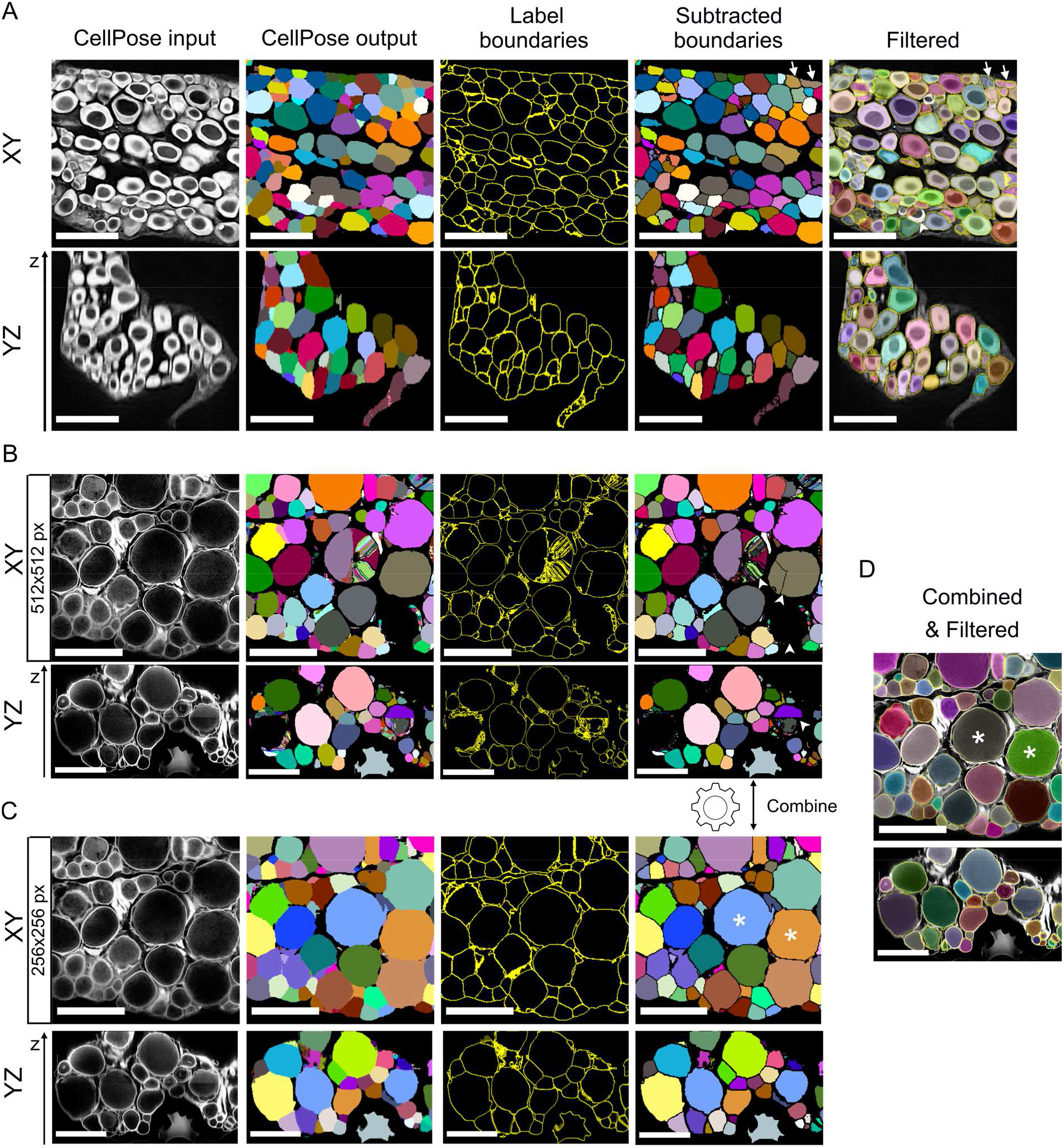
Post-processing corrections of segmented labels. (A, top) XY planes of 20 dph larvae ovary at ∼330 μm depth showing pre-treated image data, Cellpose output, labels boundaries, eroded labels (subtracted boundaries) and final results after semi-automatic label filtration in AMIRA superimposed onto image data. (A, bottom) YZ planes for qualitative assessment of segmentation along Z-axis. Segmentation results after post-processing show good accuracy and shape fitting in XY or YZ planes with only few labels fusion for smallest oocytes (arrows) or missing label (arrowhead). (B, top) XY planes of adult ovary at ∼2500 μm depth before and after Cellpose 30 segmentation. (B, bottom) YZ planes of adult ovary showing segmentation accuracy along Z axis. Largest follicles are over-segmented but other labels are correctly fitting follicles size. (C) Downscaled image data, Cellpose 30 segmentation results and post-processed images of adult data on XY plane (top) or YZ orthoslices (bottom). Large labels (stars) are combined to original scale segmentation result in (B) to replace segmentation errors of largest follicles. (D) Combined and filtered adult segmentation data superimposed onto image data. Labels show a good fit in size and shape of follicles at various sizes either on XY or YZ plane. Scale bars 100 μm (larvae panels), 1000 μm (adult panels).

### Oocyte content analyses

To assess the ovarian oocyte content at both larvae and adult stages, ovaries were imaged at each of these stages and 3D computational analyses were performed following our deep learning-based pipeline. Three-dimensional reconstructions after data pre-processing revealed the thin oval-shape of larvae ovaries oriented along the anteroposterior axis, which then evolves into a thicker rounded shape at the adult stage (Fig. 3F, 4F, 7A and 7D). Ventrally, larvae ovaries exhibited lateral folds and a marked central depression, likewise adult ovaries displayed two lateral folds as well as a ventro-median bulge, giving the ovary a wheat grain appearance. Diameters of segmented oocytes or follicles were computed, classified into different size classes and merged to the 3D ovary reconstructions (Fig. 7A’, B, D’, E). The ventral and dorsal 3D views of the larvae ovary, revealed that small oocytes were preferentially visible from the ventral views, whereas larger oocytes were only observable from dorsal views, while no obvious regionalization was observable in the adult ovary (Fig. 7B and 7E). To analyze the relative abundance of the different size classes, the developmental stage of oocytes/follicles was determined according to their diameter and as described in the oocyte developmental table of Iwamatsu *et. al*. (Iwamatsu et al., 1988). In the larvae ovary, a total of 1231 ± 182 (n=2) oocytes were detected. The mean size distribution showed a high predominance of small previtellogenic follicles ranging from 25 to 60 μm in diameter (chromatin-nucleolar stage, stage I), which suggests a synchronized oocyte growth during larval development (Fig. 7C). By contrast, all follicular developmental stages were found at the adult stage. A total of 1275 follicles were counted with a large predominance of pre-vitellogenic follicles (from stage II to IV, 50-150 μm) and of early vitellogenic follicles (stages V and VI, 150-400 μm, Fig. 7F). Proportion of follicles then progressively decrease as they progress through late vitellogenesis (stages VII and VIII, 400-800 μm). The pool of post-vitellogenic follicles (maturation stage IX, > 800 μm) is clearly distinguishable and reflects upcoming egg laying with a consistent number of about 23 follicles measuring more than 950 μm in diameter.

**Figure 7:**
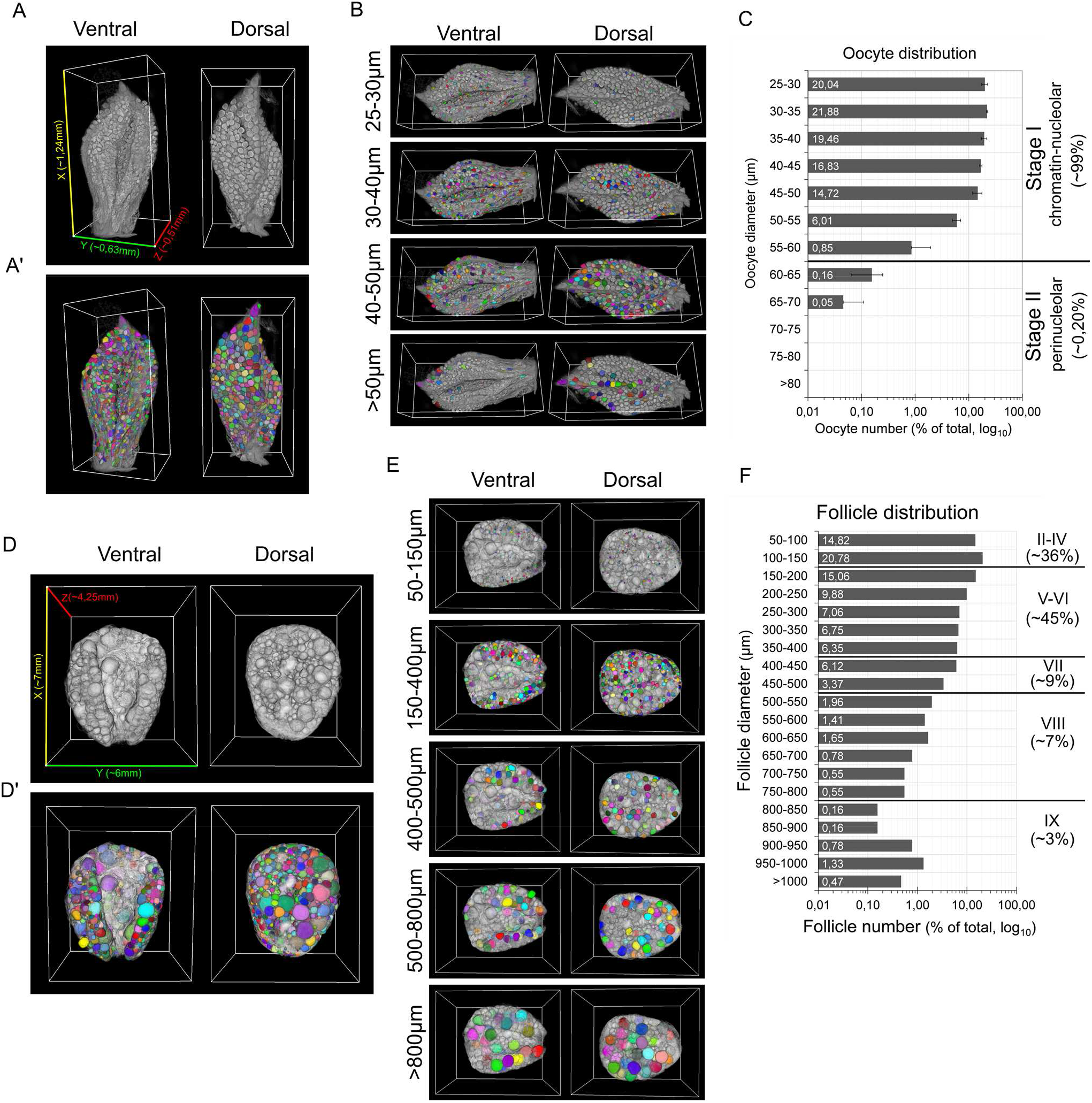
3D Qualitative spatial visualization and quantitative analysis of ovarian content. (A) 3D ventral and dorsal reconstruction views of 20 dph larvae ovary, and (A’) merged with segmented oocytes. Ovary size is approximately represented on bounding box measuring 1240 μm in length (x, yellow), 630 μm in width (y, green) and 515 μm in height (z, red). (B) Oocytes spatial distribution visualized by diameter range from ventral and dorsal side of larvae ovary. Oocytes tend to localize dorsally through their growth. (C) Oocyte distribution in entire larvae ovaries (mean +-SD, n=2) depending on their equivalent diameter. Diameter measure cut-off was applied at 25 μm. Corresponding developmental stages of previtellogenesis are indicated: stage I, chromatin-nucleolar (25-60 μm) and stage II, perinucleolar (60-90 μm). (D) 3D ventral and dorsal views of entire adult ovary after registration and reconstruction, and (D’) merged with 3D segmented follicles. Bounding box size approximates whole ovary size with an antero-posterior length of 7 mm (x, yellow), left to right width of 6 mm (y, green) and depth of 4,25 mm (z,red). (E) Follicles spatial distribution within ovary based on their equivalent diameter range. Follicle size classes show respective localization of various developmental stages, namely previtellogenesis (50-150 μm), vitellogenesis (150-800 μm) and post-vitellogenesis (>800 μm). (F) Total quantification of adult ovarian content distributed by follicular diameter. Stages of development are listed, namely previtellogenesis (II-IV, 50-150 μm), early vitellogenesis (V-VI, 150-400 μm), late vitellogenesis (VII-VIII, 400-800 μm) and post-vitellogenesis (IX, 800 μm and over). Oocyte and follicular distribution are expressed as percentage of total objects counted within the ovaries.

## DISCUSSSION

Three-dimensional imaging of whole fish ovaries typically generates large image data sets that are particularly complex to analyze. In this study, we generated two types of 3D images. On the one hand, we generated images of adult fish ovaries with low-contrast follicle outline signal at great depths, which usually greatly impairs the final segmentation efficiency, as described previously (Lesage et al., 2020). On the other hand, we generated images of larvae ovaries with a low-contrast signal inside the oocytes throughout image stacks, which makes segmentation otherwise impossible with conventional approaches. Here, we applied the generic Cellpose pre-trained algorithm that allows cell segmentation without any manual annotation and neural network training. To optimize 3D segmentation results and maximize accuracy of oocyte/follicle content analyses, Cellpose was integrated into an end-to-end analysis pipeline.

### Enhancement and homogenization of input dataset

The first part of our pipeline was dedicated to signal quality improvement in depth of raw image stacks. Such image pre-processing steps allowed improving segmentation efficiency by Cellpose. To some extent, the decrease in fluorescence level in depth on raw images should not be a major issue for predicting feature boundaries with Cellpose as it uses vector gradients representation of objects to accurately predict complex cell outlines with non-homogenous cell marker distribution (Stringer et al., 2021). However, our result indicates that the SNR is an important prerequisite for image analysis with Cellpose, in line with previous observations (Kar et al., 2021). Along with an enhanced visualization of the structures of interest across the sample, the pre-processing of 3D images therefore allows for homogenization of the data set and much more efficient 3D segmentation with Cellpose, thus increasing the reproducibility and quality of analysis.

### Improvement of Cellpose output label images

Despite its high efficiency, Cellpose led to some substantial errors, including slightly oversized or aberrant labels, and it also failed to segment oocytes of highly heterogeneous sizes. To overcome these limitations and refine labels produced by Cellpose, we performed post-segmentation corrections. The size of 3D labels was adjusted following an automated boundary subtraction strategy. Our strategy differs from other methods that use the pixel-by-pixel label erosion operation, such as in LabelsToROIs Fiji plugin designed on 2D myofiber sections, and is likely to be faster when dealing with large 3D data (Waisman et al., 2021). The combination of multiple Cellpose segmentation images, implemented with a Fiji macro “CombineLabels”, also allows identification of highly heterogeneous objects sizes, that was previously not possible with Cellpose algorithm alone. It is however worth noting that there are still few inaccuracies that could not be fixed. Under-segmentations or unsegmented objects were sometimes detected mostly with larvae image datasets. Albeit minor, these errors occur in highly oocyte-dense regions or with non-optimal signal levels. Such observation is in agreement with some studies that do not recommend Cellpose for highly overlapping masks or that describe lower accuracy with over- or underexposed images (Kar et al., 2021). This could be attributed to the 2D averaging process for the 3D Cellpose extension that may have lower accuracy than a model trained with 3D data, especially for highly dense regions (Lalit et al.,2022; Stringer et al., 2021). Obviously, one can assume that better accuracy could be achieved by using a dedicated specialized DL model, and in particular with 3D trained model on our data, as shown by D.Eschweiler et *al* (Eschweiler et al., 2022). It would thus be interesting in the future to use our segmentation results for Cellpose algorithm fine-tuning. This could indeed limit the need for image pre-processing as well as post-processing corrections of segmentation results. But in this case, we would somewhat lose the advantage of versatile generalist models like Cellpose and different models would have to be trained for each type of data. Another solution could therefore be to improve the input images quality, by using a suitable oocyte marker to avoid sharp signal enhancement and posssibly in combination with a membrane marker for better boundary discrimination. Alternatively, and in absence of such specific staining, another denoising process, either trained in three dimensions, with noisy/non-noisy paired images (CARE) or combining deconvolution process (DecoNoising), could also help objects recognition accuracy (Weigert et al., 2018; Goncharova et al., 2020).

### An accurate and comprehensive content analysis of larvae and adult medaka ovaries

Implementation of Cellpose for oocytes/follicle 3D segmentation eventually enabled unbiased, reproductible and comprehensive studies for meaningful biological information, which offers great possibilities for a complete description of fish ovarian growth and development. From a morphological point of view, we could clearly distinguish the oval shape of the ovary thickening over time and shaping a bulge in the ventro-median position that connects the mesentery and attaches to the gut (Iwamatsu, 2015; Lesage et al., 2020). *In situ* follicular size measurements by our 3D imaging and DL-based segmentation approaches allowed producing size distribution profiles for both larvae and adult ovaries. Our results are consistent with those obtained previously from dissociated follicles measured manually for the larvae ovary (Iwamatsu, 2015) or semi-automatically from 3D images by classical watershed segmentation approaches for the adult ovary as shown in our previous study (Lesage et al., 2020). However, greater confidence can be attributed to the present study, particularly for the pre- and post-vitellogenic stages in the adult ovary for which we achieved fewer segmentation errors. In general, we also achieved a better estimation of follicle size due to the accurate shape detection enabled by the Cellpose algorithm. Interestingly, we also noticed that the spatial distribution of oocytes between 30 and 70 μm in diameter tended to be regionalized along the ventro-dorsal axis in the larvae ovary, suggesting an oriented follicular growth through this axis in consistency with observation of Nakamura *et al*. (Nakamura, 2018). In the future, the ovarian morphogenesis and spatial organization of follicles according to their size should however be further characterized during the ovarian development by using refined 3D spatial analysis approaches.

## Conclusion

Overall, the use of the generic Cellpose algorithm has been successful for 3D ovary images and has allowed ovarian segmentation of unprecedented quality. Cellpose significantly accelerated and improved the efficiency and the quality of ovarian follicles 3D segmentation in adults, leading to an accurate count and measurement of all oocyte diameters. Even more remarkably, this generalist model also allowed the successful segmentation of images of larvae ovaries with weak fluorescent signal, otherwise not exploitable with conventional methods, and quite certainly even after image pre-processing. This possibility challenges the dogma that a good raw image is necessary for an accurate object segmentation and thus significantly increases further analysis opportunities. Furthermore, thanks to its ease of use, implementation of Cellpose avoids the tedious and complex step of setting up an AI segmentation method and is therefore largely accessible to non-specialist biologists with limited coding and hardware knowledge. In the deep learning era, it is thus now clearly possible to apply such a cutting-edge technology for tissue 3D phenotyping with relative ease. To our knowledge, our pipeline is the first application using developer-to-user deep learning solutions for 3D image analysis of the ovary in vertebrates, thus opening the way for further innovative in-depth morphometric studies within the framework of developmental or toxicological studies.

## ACKNOWLEDGMENTS

We thank the INRAE ISC-LPGP fish facility staff and especially Amélie Patinote and Guillaume Gourmelen for fish rearing and husbandry.

## ADDITIONAL INFORMATION AND DECLARATIONS

### Author Contributions

M.L. performed the experiments, the computational analyses and wrote the manuscript. J.B. participated to the setup of the computation bio-image analyses. M.T. participated in the setup of the clearing protocol and to the image acquisition. T.P. participated in the choice and implementation of the tool for the 3D registration of adult ovary images. V.T. conceived the study, participated in data analyses and manuscript writing. All authors reviewed drafts of the article and approved the final manuscript.

